# Evolutionary Flexibility of Ribosome Biogenesis in Bacteria

**DOI:** 10.1101/2024.11.29.625318

**Authors:** Kazuaki Amikura, Shun’ichi Ishii, Yoshihiro Shimizu, Shino Suzuki

## Abstract

Ribosomes are essential for protein synthesis and require ribosome biogenesis factors (RBFs) for assembly. To uncover the evolutionary diversity of ribosome biogenesis, we analyzed over 30,000 bacterial genomes and revealed that Candidate Phyla Radiation (CPR), also known as the phylum Patescibacteria, characterized by reduced genomes and smaller ribosomes, has about half the average number of RBFs compared with non-CPR bacteria. Notably, key RBFs such as der, obgE, and rbfA, considered indispensable, are conserved in only around 20%–70% of CPR genomes. Since such repertoires were not observed in reduced genomes of other phyla, CPR presumably diverged early in bacterial evolution. We further confirmed that ribosomal structural changes correlate with reduced RBFs, evidencing co-evolution between RBFs and the ribosome. These findings suggest that ribosomal biogenesis is more flexible than recognized, and the small cell and genome sizes of CPR bacteria and their early divergence may influence the unusual repertoires of RBFs.

**Teaser:** Ribosome biogenesis in CPR bacteria was unexpectedly flexible, challenging traditional views of this essential process in evolution.

## Introduction

Ribosomes serve as essential macromolecular machines that synthesize proteins within cells and consist of ribosomal RNAs (rRNAs) and numerous ribosomal proteins (RPs). The synthesis of new ribosomes is a highly complex and energy-intensive process, crucial for maintaining cellular function and life (*1–4*). This process requires the coordinated synthesis, processing, and modification of rRNA and RPs, along with intricate folding and hierarchical assembly of these components. Ribosome biogenesis factors (RBFs), including GTPases, ribonucleases, helicases, modification enzymes, and chaperones, play crucial roles in ensuring the accurate and efficient assembly, folding, processing, and modification of ribosomal components within the cell (Fig. 1A). While significant advances have been made in understanding ribosome biogenesis, the precise mechanisms enabling cells to rapidly and accurately synthesize ribosomes, as well as the evolutionary origins of these highly conserved systems, remain unclear. Most bacteria possess approximately 50 RBF genes (*5–7*), with approximately a dozen being essential for the survival of *Escherichia coli* (*8*). The Genome Taxonomy Database (GTDB), a comprehensive resource that provides a standardized taxonomy for prokaryotes based on genome phylogeny, includes 21 RBFs in the core gene sets for phylogenetic analyses, emphasizing their evolutionary significance (*9*, *10*). Furthermore, 15 RBFs belong to protein families that trace back to the Last Bacterial Common Ancestor (LBCA) (*11*) (Supplementary Table 1), linking RBFs to broader biological evolution.

**Fig. 1.**
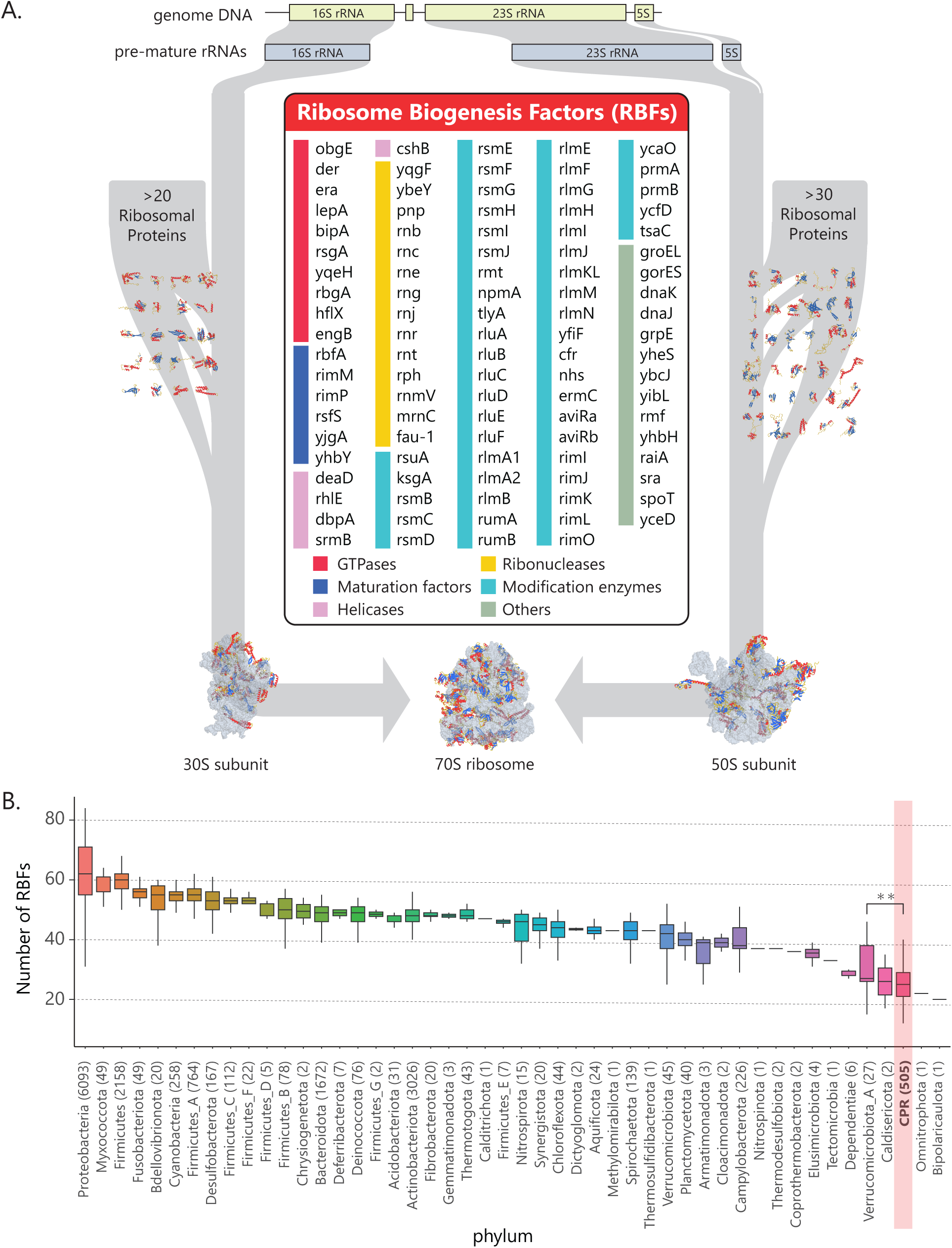
Ribosome biogenesis processes and sets of biogenesis factors (BFs) in bacteria. (**A**) Schematic view of multi-step ribosome biogenesis including the list of ribosome biogenesis factors and tertiary structure of ribosomal proteins, 70S, 50S and 30S subunits (PDB code: 4YBB, 5UYN, 6ENF, and 1RQU). (**B**) Boxplot representation of the number of BFs across different phyla in bacteria. Firmicutes are divided into seven groups in the AnnoTree database.

While rRNAs, RPs, and RBFs are generally highly conserved across the bacterial domain, recent studies have reported that Candidate Phyla Radiation (CPR) bacteria, a taxonomic group of bacteria characterized by small genome sizes ranging from 0.5 to 1.5 Mbp and small cell sizes of 0.2 - 0.3 µm, exhibit unusual characteristics related to the ribosome (*12–14*). CPR bacteria commonly contain introns in their rRNA and have atypical ribosome structures, with bL30 absent across all CPR lineages and ribosomal proteins such as uL1 and bL9 absent in a lineage-specific manner (*12*, *15*). Recent study reported that several ribosomal proteins in the large subunit also absent in a lineage-specific manner (*16*). Additionally, the gene *der*, which belongs to the RBFs and generally remains conserved across all bacteria, appears absent in some CPR classes (*12*). Although researchers continue to discuss the taxonomic position of CPR bacteria (*17*), CPR is mostly recognized as a deeply branched lineage in the bacterial tree of life.

The unusual gene distributions of RPs and the phylogenetic uncertainty observed in the CPR bacteria offer unique and intriguing opportunities to explore the diversity and evolution of ribosome biogenesis processes as well as the CPR phyla. In this study, we conducted a comprehensive analysis of RBFs across the domain Bacteria using high-quality bacterial genome sets, with a special focus on CPR bacteria. By analyzing high-quality bacterial genome sets, we aimed to reveal the diversity and evolutionary flexibility of ribosome biogenesis in this unique group, providing insights into the evolutionary trajectory of RBFs and CPR phyla.

## Results

### Lists of ribosome biogenesis factors and high-quality bacterial genomes

From the Kyoto Encyclopedia of Genes and Genomes (KEGG) database, we used all 83 RBFs that were classified as a ribosome biogenesis factor of prokaryotic types and manually curated an additional 17 RBFs based on previous studies, including *lepA, bipA, engB, rhlE, pnp, rnb, rnj, rnr, rnt, rph, rnmV, groEL, groES, dnaK, dnaJ, grpE,* and *yheS* (Fig. 1A, Supplementary Table 1). A total of 100 RBF genes were identified and categorized into six types: GTPases, maturation factors, ribonucleases, helicases, and modification enzymes, and others.

To comprehensively analyze the gene distribution of RBFs in bacteria with high-quality and well-annotated genomes, we initially examined 15,274 bacterial genomes registered in RefSeq, selected from over 30,000 bacterial genome sets derived from AnnoTree version 1.2 (http://annotree.uwaterloo.ca). AnnoTree enables interactive exploration of the taxonomic distribution of genes across comprehensive bacterial species (*18*, *19*); However, the limited availability of only five genomes of CPR bacteria in the RefSeq database poses challenges in accurately assessing the gene distribution of CPR bacteria. To overcome this limitation, we collected high-quality metagenome-assembled genomes (MAGs) of CPR bacteria from GenBank, and assembled additional MAGs in our laboratory (See Material and Methods). In total, 505 high-quality MAGs of CPR bacteria were obtained with a median completeness of 97.4%, as calculated using CheckM2 (*20*). Nearly full-length 16S, 23S, and 5S rRNAs were identified in all MAGs of CPR bacteria, with the median length of 16S being 1390 bases and that of 23S being 2384 bases. The 3ʹ-terminus of 16S rRNA was often not fully annotated. Introns were frequently observed in the 16S and 23S rRNA. Furthermore, we added six high-quality MAGs from the phylum Dependentiae, as atypical RPs have been reported in Dependentiae, along with their protein family content distribution (*12*, *21*). RBF gene sets from 505 CPR and 15,274 non-CPR bacterial genomes were prepared for comprehensive analysis.

### Smaller number of ribosome biogenesis factors in CPR bacteria

Notably, the median number of RBFs identified in the genomes of CPR bacteria was 25, which is nearly half the average number of RBFs in non-CPR bacteria (Fig. 1B, Fig. 2B). Given that their ribosomal structures are not significantly different, with similar values for ribosomal RNA length and the median number of ribosomal proteins (50 in CPR bacteria and 53 in non-CPR bacteria), the observed loss of RBFs in CPR bacteria was unexpected (Fig. 2A, Supplementary Fig. 1). Principal component analysis (PCA) based on the profiles of RBFs revealed clear differences in the distribution of RBFs between CPR and non-CPR bacteria (Fig. 2C). Although the phyla Bipolaricaulota and Omnitrophota had fewer RBFs than CPR bacteria, further consideration is required due to the limited availability of genomes for these phyla (Fig. 1B).

**Fig. 2.**
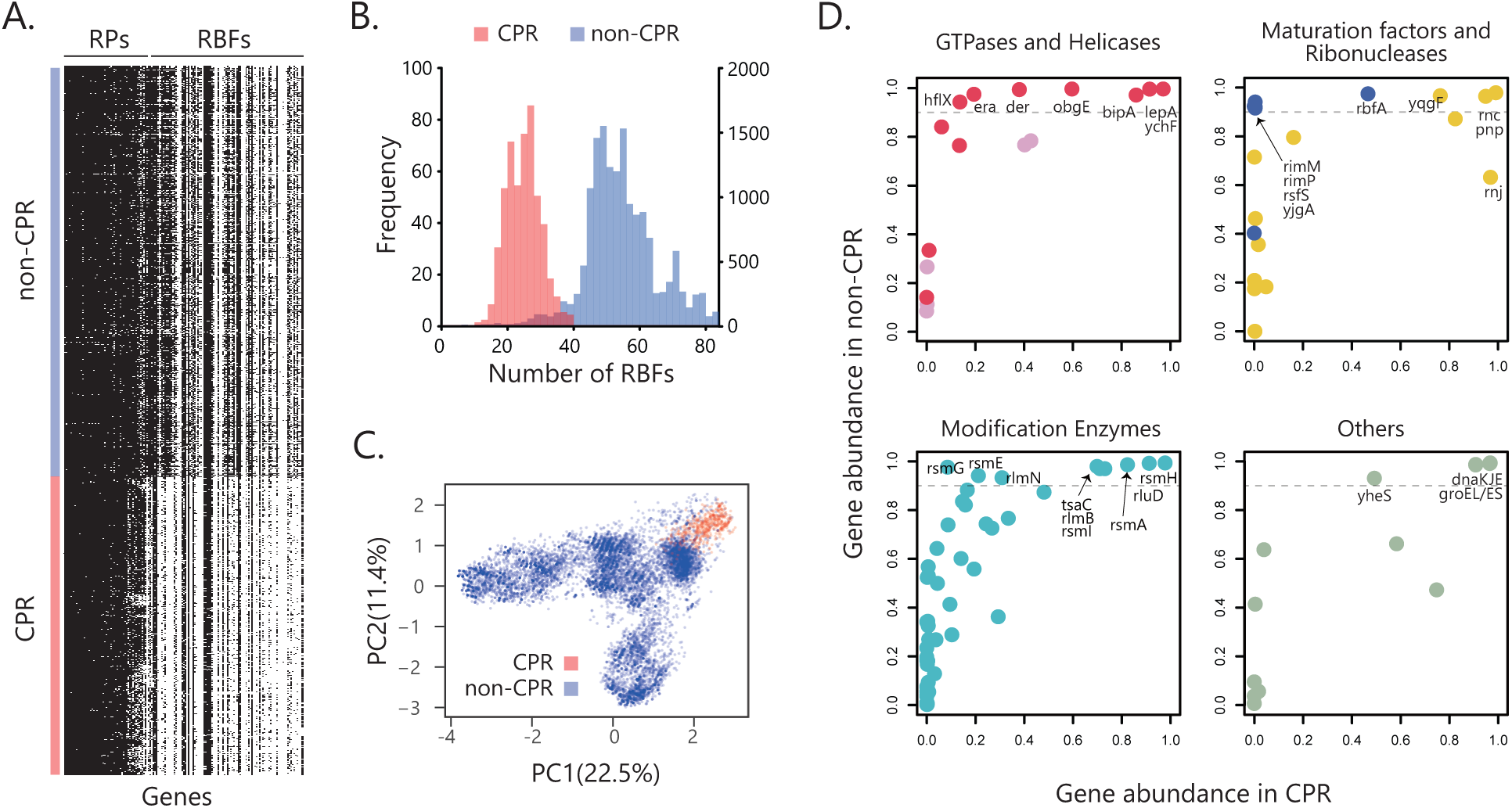
Analysis of gene distributions of biogenesis factors (BFs) between Candidate Phyla Radiation (CPR) and non-CPR bacteria. (**A**) The gene distribution of ribosomal proteins (RPs) and BFs (columns) in CPR bacteria and non-CPR bacterial genomes (rows) that show 694 genomes, one representative from each family, out of non-CPR genomes sets in this study. (**B**) The histogram represents the distribution of the number of biogenesis factors or modification enzymes in a genome from CPR bacteria (Red, Left vertical axis) and non-CPR (Blue, Right vertical axis). (**C**) Principal component analysis (PCA) was performed on the presence or absence of RBF sets among 15,779 genomes. (**D**) A comparison of gene conservation from the constructed genome sets in this study between non-CPR bacteria and CPR bacteria (Supplementary Table 3). Gray dot lines show that the conservation ratio in non-CPR bacteria is 0.9.

Among the 30 RBFs that were highly conserved within the non-CPR bacteria (conservation ratio >90%), 14 RBFs showed a lower conservation ratio (ranging from 10% to 90%) within the CPR bacteria (Fig. 2D, Supplementary Table 2). The RBF genes *obgE* and *der*, both of which are essential for the maturation of the 50S ribosomal subunit (*22–25*) and whose deletion is lethal in *E. coli* (*8*), were conserved in over 99% of non-CPR bacteria. However, they were significantly less conserved in the CPR bacteria, with conservation ratios of 60% and 38%, respectively. The other 12 RBFs that were less conserved in CPR bacteria included GTPases *era*, *bipA*, and *hflX*, maturation factor *rbfA*, ribonuclease *yqgF*, and modification enzymes *rlmN*, *rsmE*, *rsmI*, *rsmA*, *tsaC*, and *rlmB*, and ATP-binding protein *yheS*, all of which were highly conserved in non-CPR bacteria. *rsmG* exhibited a high conservation ratio (> 90%) in non-CPR bacteria, in contrast to the low conservation ratio (<10%) observed in CPR bacteria. Similarly, *rimM*, *yjgA*, *rsfS*, and *rimP* were nearly absent in the CPR bacteria, in contrast to the high conservation rates (> 90%) observed in the non-CPR bacteria.

### Class-specific gene distribution of RBFs in CPR bacteria

The phylogenetic distribution of the less conserved (greater than 10% and less than 90%) RBF and RP genes in CPR bacteria was analyzed to explore the evolutionary trajectory of these genes within CPR bacteria. These genes included 14 RBF genes encoding GTPases (*der, obgE, bipA, hflX*, and *era*), a maturation factor (*rbfA*), a ribonuclease (*yqgF*), modification enzymes (*rlmN*, *rsmE*, *rsmI*, *rsmA*, *tsaC*, and *rlmB*), an ATP-binding protein *yheS*, and two RPs (uL1 and bL9; Fig. 2D). The results revealed that four of the 14 RBF genes, *der, obgE, era*, and *rbfA*, and 2 RPs exhibited class-specific distributions, whereas the remaining genes were broadly distributed across CPR bacteria. Detailed analysis of *der* and *obgE*, both of which function in the later stage of 50S maturation, indicated that the two exhibit a complementary relationship within the CPR bacteria. Although most CPR bacteria maintain *obgE*, ABY1 lacks *obgE* but maintain*s der* (Fig. 3A,B). Conversely, four classes in CPR (Paceibacteria, Dojkabacteria, WWE3, and Microgenomatia) lacked *der* but retained *obgE*. Similarly, both *era* and *rbfA* that are involved in 30S rRNA biogenesis, showed the complementary trends among the classes ABY1 and Paceibacteria (Fig. 3B). The classes Dojkabacteria, WWE3, and Saccharimonadia, which are phylogenetically close within the CPR, almost lacked both *era* and *rbfA*.

**Fig. 3.**
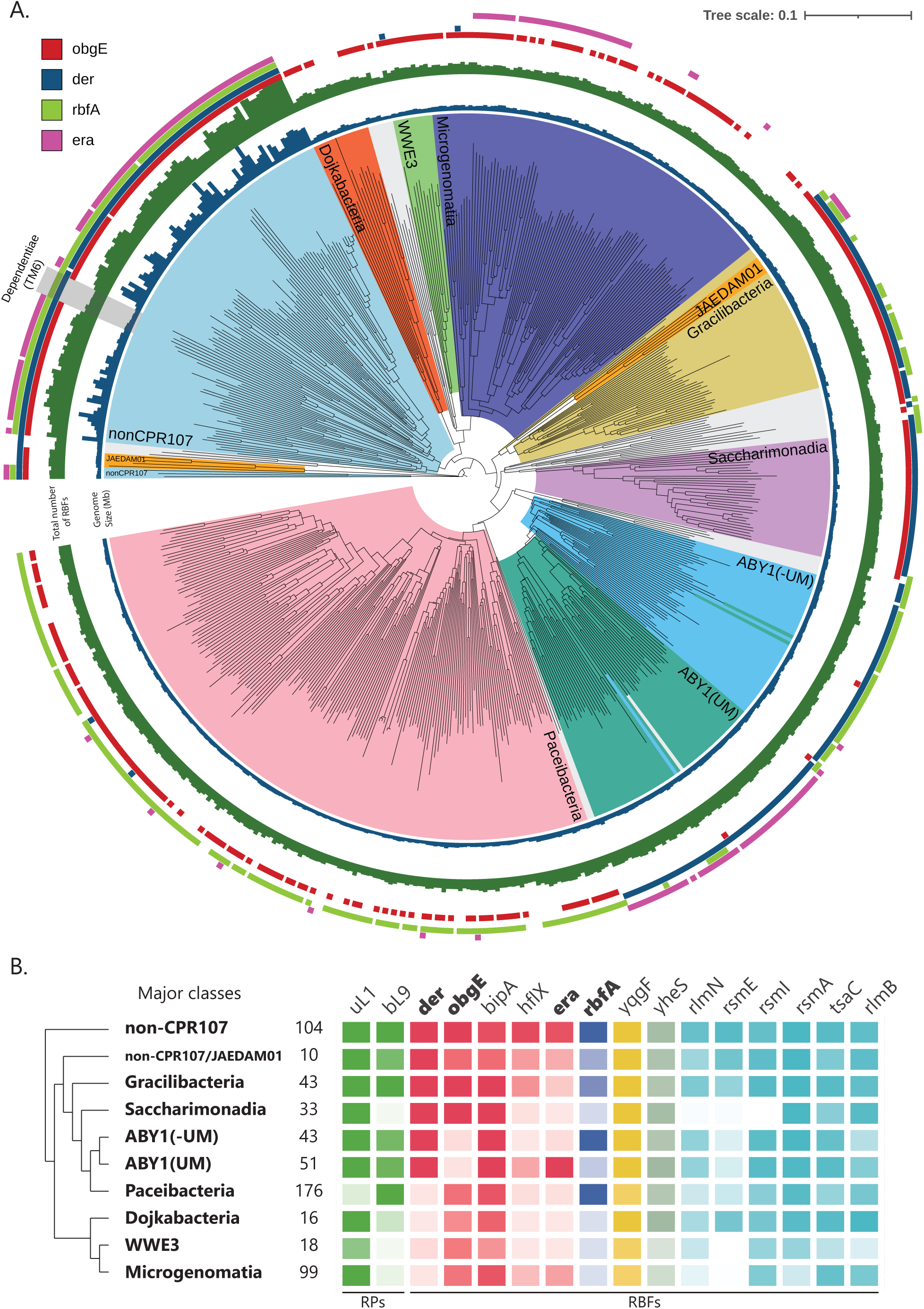
Distribution of the ribosome biogenesis factors (RBFs) in Candidate Phyla Radiation (CPR) bacteria. (**A**) The tree was constructed by the concatenated alignment fasta file made by checkM. non-CPR107 including Dependentiae selected from representative species from each phylum (Supplementary information). Leaves are colored according to classes of CPR, with the top eight classes containing the highest number of species highlighted in distinct colors. Classes with fewer species are shown in gray, and species names are not labeled on the tree (Supplementary Information). ABY1(UM) indicates Uhrbacteria or Magasanikbacteria in NCBI taxonomy. Blue bars indicate the genome size. Green bars indicate the number of biogenesis factors (BFs). The presence of 4 RBFs is marked by a colored square (inner to outer arcs, respectively). (**B**) The conservation within each clade of genes. The intensity of color reflects the level of conservation, with darker shades indicating high conservation. The number represents the number of species in the clade.

### der as a key factor for the unusual distribution of RBFs and RPs

Another PCA analysis was performed only for CPR bacteria based on the presence or absence of RBF and RP genes to examine the gene distribution pattern related to ribosome biogenesis of CPR bacteria (Fig. 4A). The analysis revealed that closely related classes within CPR bacteria exhibited overlapping confidence ellipsoids, whereas distantly related classes showed no overlap. For example, classes Saccharimonadia and Paceibacteria were taxonomically distinct and displayed unique distribution patterns (Fig. 3B). Further, the factor loading analysis indicated a substantial influence of *der*, *rbfA*, *uL1*, and *bL9* (Fig. 4B). These genes represented the top four contributors to the principal components identified through PCA, with *der* mainly contributing to PC1 and *rbfA*, uL1, and bL9 contributing to PC2.

**Fig. 4.**
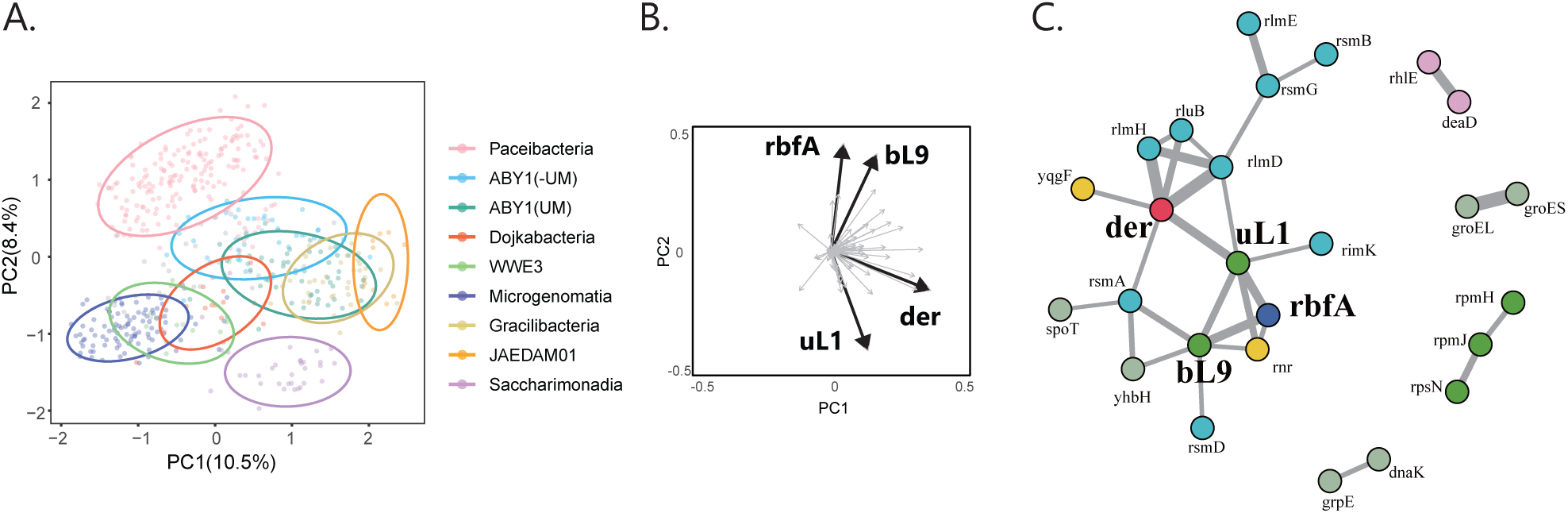
Principal component analysis (PCA) and dependency network. (**A**) PCA was performed using ribosomal proteins (RPs) and ribosome biogenesis factors (RBFs). The ellipses in the plot represent 90% confidence ellipse intervals. Principal component loadings are shown on the adjacent plot. The top four factors with high principal component loadings are indicated by bold arrows. (**B**) The network represents the dependency based on the mutual information. The width of the edge is proportional to the value of mutual information. The yellow node represents biogenesis factors (BFs). The blue node represents RPs.

To gain a comprehensive understanding of the mechanisms underlying the unique RBF/RP distribution patterns in the CPR bacteria, a mutual information analysis was performed (Fig. 4C). This quantitative approach calculates the dependency between two genes via protein-protein interactions and suggests their potential involvement in similar biological processes (*26*). Among 100 RBFs and 54 RPs, 11,781 potential interactions were evaluated, and 30 substantial dependencies were revealed with a mutual information value of 0.1 or higher (Supplementary Fig. 2). The connection between the chaperone *groEL* and its cofactor *groES* showed the highest value of mutual information at 0.336, indicating co-occurrence or co-absence in CPR bacteria (Supplementary Table 3). Notably, *der* exhibited the largest number of connections and the highest cumulative edge weight among all RBFs and RPs (Supplementary Table 4), indicating that *der* plays a pivotal role in ribosome biogenesis in CPR bacteria via its association with the ribosomal protein *uL1*, ribonuclease *yqgF*, and several modification enzymes, including *rlmD*, *rlmH*, *rluB*, and *rsmA* (Fig. 4B). The significance of *der* in the network remained consistent, even when the threshold for defining the dependency of the mutual information analysis varied (Supplementary Table 4, Supplementary Fig. 3). These results indicated that *der* is a key factor in the unique gene distribution patterns of RPs and RBFs in CPR bacteria and likely plays a pivotal role in ribosome biogenesis.

### rRNA structure co-evolved with ribosomal proteins and biogenesis factors

We further analyzed rRNA secondary structures to understand the correlation between the distribution patterns of RPs and RBFs. Since the rRNA length in CPR bacteria is generally shorter than that in non-CPR bacteria (*16*), we analyzed 50, 5, and 100 known helices of 16S, 5S, and 23S rRNA genes, respectively, of domain bacteria and compared the length of the helices to those of CPR bacteria. The analysis showed that seven helices of 16S, 11 helices of 23S, and one helix of 5S rRNA in CPR bacteria were shorter than those in the seed sequences, which were a curated set of representative sequences for each rRNA obtained from the Rfam database (Supplementary Fig. 4). We also observed that helices H12, H58, and H78, were notably shorter and distributed in a class-specific manner (Fig. 5).

**Fig. 5.**
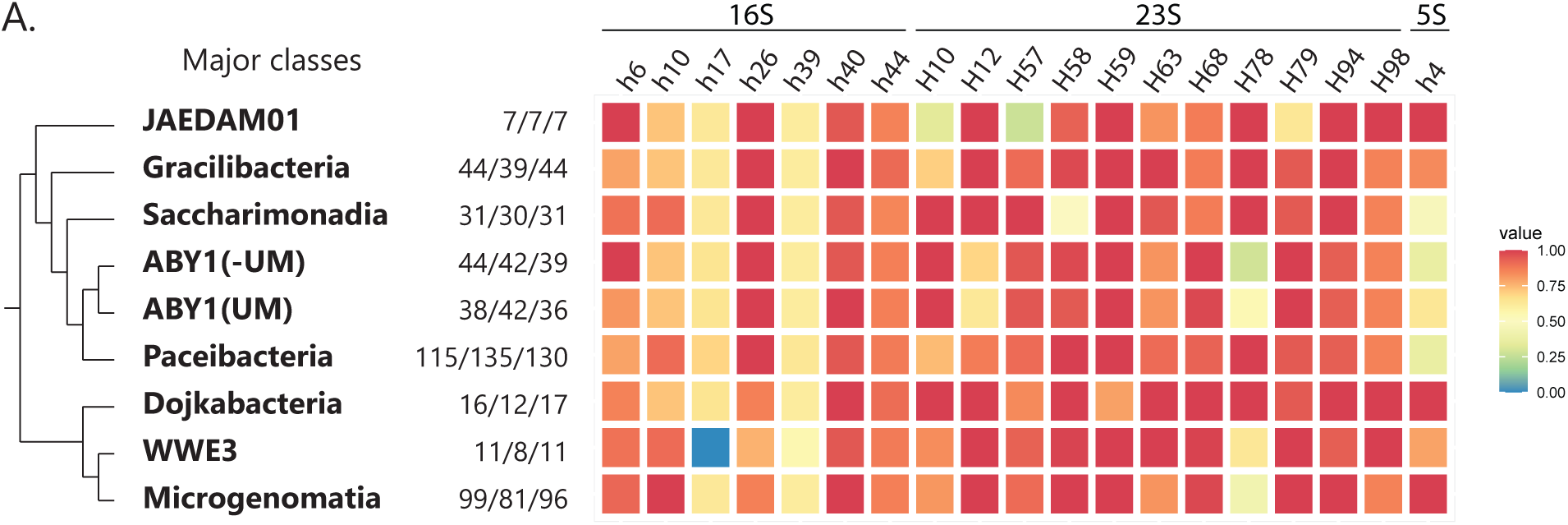
Truncated rRNA structures in CPR bacteria. The heatmap represents the median length of helices. The numbers next to the heatmap shows the number of sequences from 16S/23S/5S rRNA.

To analyze the class-specific evolutionary process among rRNA structure, RPs, and RBFs, we focused on H78 of 23S rRNA, because they are proximal to *der* and uL1, and also showed class-specific distributions in CPR bacteria (Fig. 5). During 50S subunit biogenesis, GTPase Der and ribosomal protein uL1 interact with the immature 50S subunit, and the N-terminus of uL1 interacts with H78 (*25*) (Fig. 6A, Supplementary Fig. 5) The L1 stalk region, formed by H76, H77, H78, and uL1, plays a critical role in translation with dynamic structural movements to facilitate the release of tRNA from the ribosome (*27*). The presence or absence of these three components (Der, H78, and uL1) was determined based on CPR phylogenies, which showed that the classes ABY1, WWE3, and Microgenomatia lacked H78 (Fig. 6B). The length and helical structure of H78 gradually changed during diversification (Fig. 6C, Supplementary Fig. 6). We found that the internal loop, which is a common structure in non-CPR bacteria, of H78 disappeared in three classes of CPR bacteria (ABY1, WWE3, and Microgenomatia) and had the same sequence length and structure as those in non-CPR bacteria (Fig. 6B, Supplementary Information). The shorter H78 species were divided into two categories. One had a stem length of 2–3 base pairs, whereas the other had a stem length of 6– 8 base pairs (Fig. 6B). In the same clade, H78 of Paceibacteria and ABY1 showed gradual changes (Supplementary Fig. 6), whereas ABY1(UM) only contained species with a shortened H78 length. As ABY1(UM) branched from ABY1, only species with a shorter H78 were retained. Similarly, changes in H78 length were observed in Dojkabacteria, WWE3, and Microgenomatia (Supplementary Fig. 6).

**Fig. 6.**
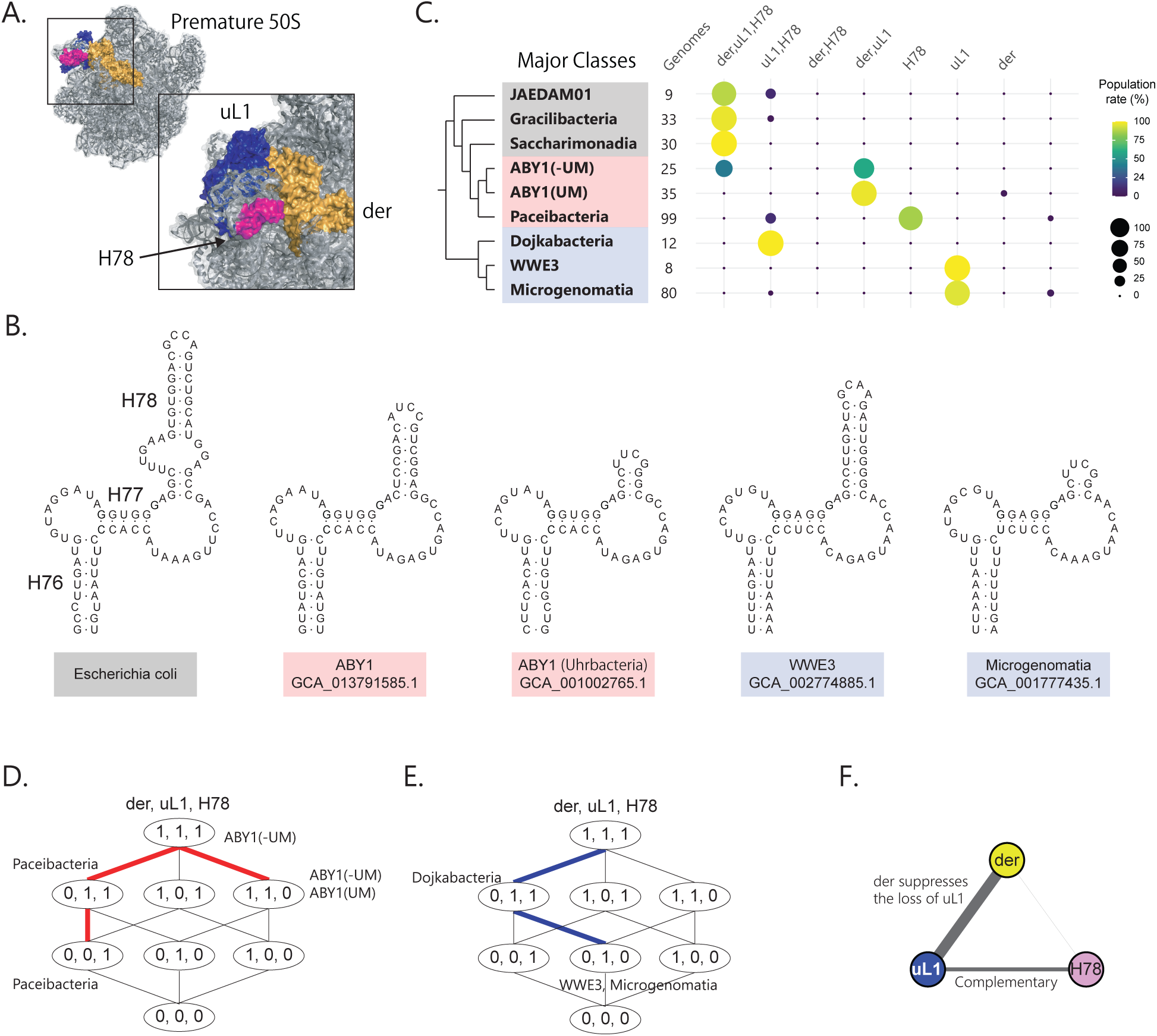
Co-evolutionary pathway of ribosome structure and ribosome biogenesis. (**A**) Der bound premature 50S subunits: PDB 3J8G. (**B**) Representative secondary structures of H78 in Candidate Phyla Radiation (CPR) bacteria and *Escherichia coli*. (**C**) Dot plot of the presence or absence of *der*, *uL1*, and *H78* within each class. Each combination of genes is quantified and its proportion relative to the total count within the classes is depicted by the size and color of circles. (**D**) Co-evolution pathway of uL1, H78, and Der. The number within the ellipse represents the combination of the presence (1) or absence (0) of der, uL1, and H78. Red line indicates the path in the clade consisting of ABY1 and Paceibacteria. The blue line indicates the path in the clade consisting of Dojkabacteria, WWE3, and Microgenomatia. (**E**) Dependency between der, uL1, and H78. The width of the edge is proportional to the value of mutual information.

To investigate whether *der* and uL1 genes co-evolved along with structural changes in H78, the presence or absence of these genes was determined based on CPR phylogenies (Fig. 6C). Two distinct evolutionary trajectories were revealed for these combinations within CPR bacteria (Fig. 6D). Although ABY1 and Paceibacteria exhibited a strong correlation in terms of phylogenomic position, *der*/uL1/H78 genes in these classes exhibited disparate evolutionary paths (Fig. 6D). ABY1 gradually loses H78 and maintains *der* and uL1, whereas Paceibacteria maintains H78 but lacks uL1 and *der*. Interestingly, in the Dojkabacteria, WWE3, and Microgenomatia clade, Dojkabacteria first lost der, but instead of losing uL1, as in Paceibacteria, they lost H78 next (Fig. 6E).

Calculations using mutual information based on the *der*/uL1/H78 profiles from 505 CPR genomes indicated a high dependency between *der* and uL1, and the lowest dependency was observed between H78 and *der*, suggesting that *der* strongly suppresses the loss of uL1, whereas H78 and uL1 are in a complementary relationship (Fig. 6F). These results indicate that these three factors have co-evolved and show a diversity of evolutionary trajectories through the evolution of ribosome biogenesis and structures.

### Comparison between symbiotic/parasitic bacteria and CPR bacteria

Given that reduced genomes are also observed in symbiotic and parasitic bacteria because of relatively recent genome streamlining, we compared the RBF distribution profiles of these bacteria and CPR bacteria to examine whether the unusual distribution of RBFs is a common feature of reduced genomes. We manually selected 438 known symbiotic and parasitic bacteria from a wide range of taxa (Supplementary Fig. 7A). These included intracellular endosymbionts of insects, such as *Orientia tsutsugamushi*, *Chlamydophila*, *Mycoplasma,* and *Phytoplasma* which infect blood, animals, and plants, respectively. Compared to the symbiotic, parasitic and CPR bacteria, the average genome size of CPR bacteria was approximately 20% smaller (Fig. 7A), and the number of RBFs per genome were >30% lower (Fig. 7B). A PCA of the entire dataset, as well as subsets containing only CPR bacteria and symbiotic and parasitic bacteria, revealed that non-CPR bacteria were divided into two distinct clades, each consisting primarily of Terrabacteria and Gracilicutes groups (Supplementary Fig. 7B). Symbiotic and parasitic bacteria showed greater diversity in the distribution of RBFs than CPR bacteria (Fig. 7C). Further detailed analyses revealed that over 99% of the symbiotic and parasitic bacteria conserved *obgE*, whereas only 60% of the CPR bacteria conserved this gene. This trend was also observed in *der*, *rbfA*, and *era* (Fig 7D). Conversely, the RBF genes *spoT*, *yhbH*, and *yheS* were less-frequently conserved in symbiotic/parasitic bacteria, but highly conserved in CPR bacteria. These findings indicated that the number of RBFs was generally lower in the reduced genomes of both CPR bacteria and symbiotic/parasitic bacteria, but the distribution patterns differed between the two groups. The *obgE, der, rbfA*, and *era* genes were highly conserved in symbiotic/parasitic bacteria but less conserved in CPR bacteria.

**Fig. 7.**
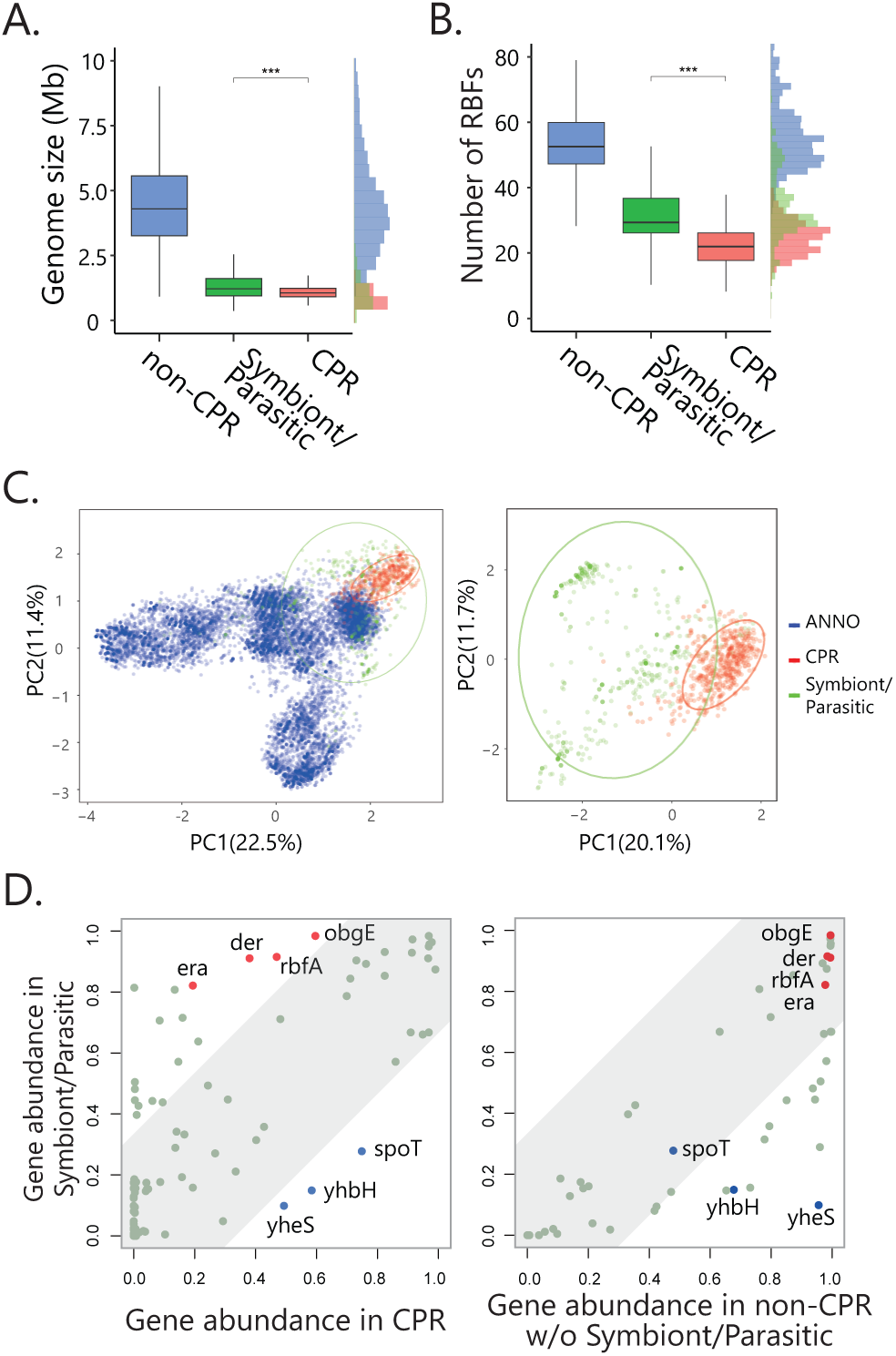
Comparison between symbiotic bacteria and Candidate Phyla Radiation (CPR) bacteria. Boxplot comparing genome size (A) and the number of biogenesis factors (BFs; **B**). (**C**) The histogram represents the distribution of the number of BFs or modification enzymes in a genome from CPR bacteria (Red, Left vertical axis), non-CPR (Blue, Right vertical axis), and symbiotic bacteria (Green, Left vertical axis).

### Gain and loss analysis of *der*, *obgE*, *era* and *rbfA* in CPR bacteria

Phylogenetic trees of *obgE, der, rbfA*, and *era* in CPR bacteria tended to form distinct clusters separate from non-CPR bacteria, and these trees were largely consistent with the tree of bacterial life determined by the GTDB (Fig. 8). However, some exceptions were observed, including gene transfers from Gracilibacteria and JAEDAM01 to non-CPR bacteria, as well as the gene transfer of *obgE* from Alphaproteobacteria to CPR clades. To understand the relationship between the phylogeny of CPR bacteria and the gene distribution of *obgE, der, rbfA*, and *era,* gene gain and loss analyses of the four RBF genes were conducted (Fig. 9). Consequently, the closely related groups of classes Dojkbacteria, WWE3, and Microgenomatia maintained *obgE* but lost the remaining three, whereas part of Microgenomatia gained *era*. The classes Paceibacteria, ABY1, Saccharimonadia, and Gracillibacteria have different combinations of RBFs, but mostly do not have *era* and the ABY1 (UM) gained *era* during the later stages of evolution. These results suggest that the current set of RBFs was largely established before each class was formed, and that the classes Dojkbacteria, WWE3, and Microgenomatia may have diverged from a common ancestor.

**Fig. 8.**
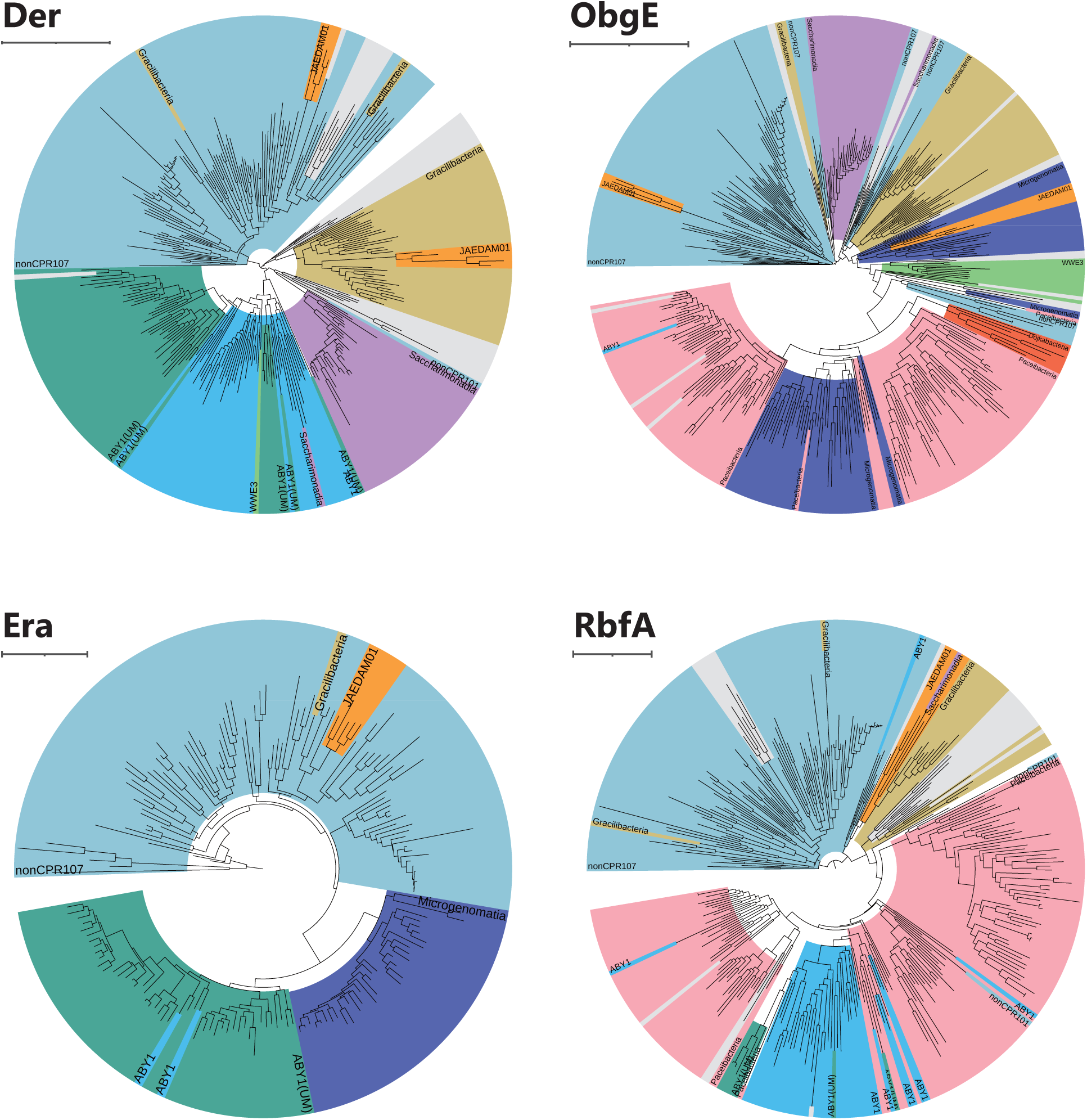
Phylogenetic trees of four ribosome biogenesis factors (RBFs) The tree was constructed using IQ-TREE and rendered using iToL.

**Fig. 9.**
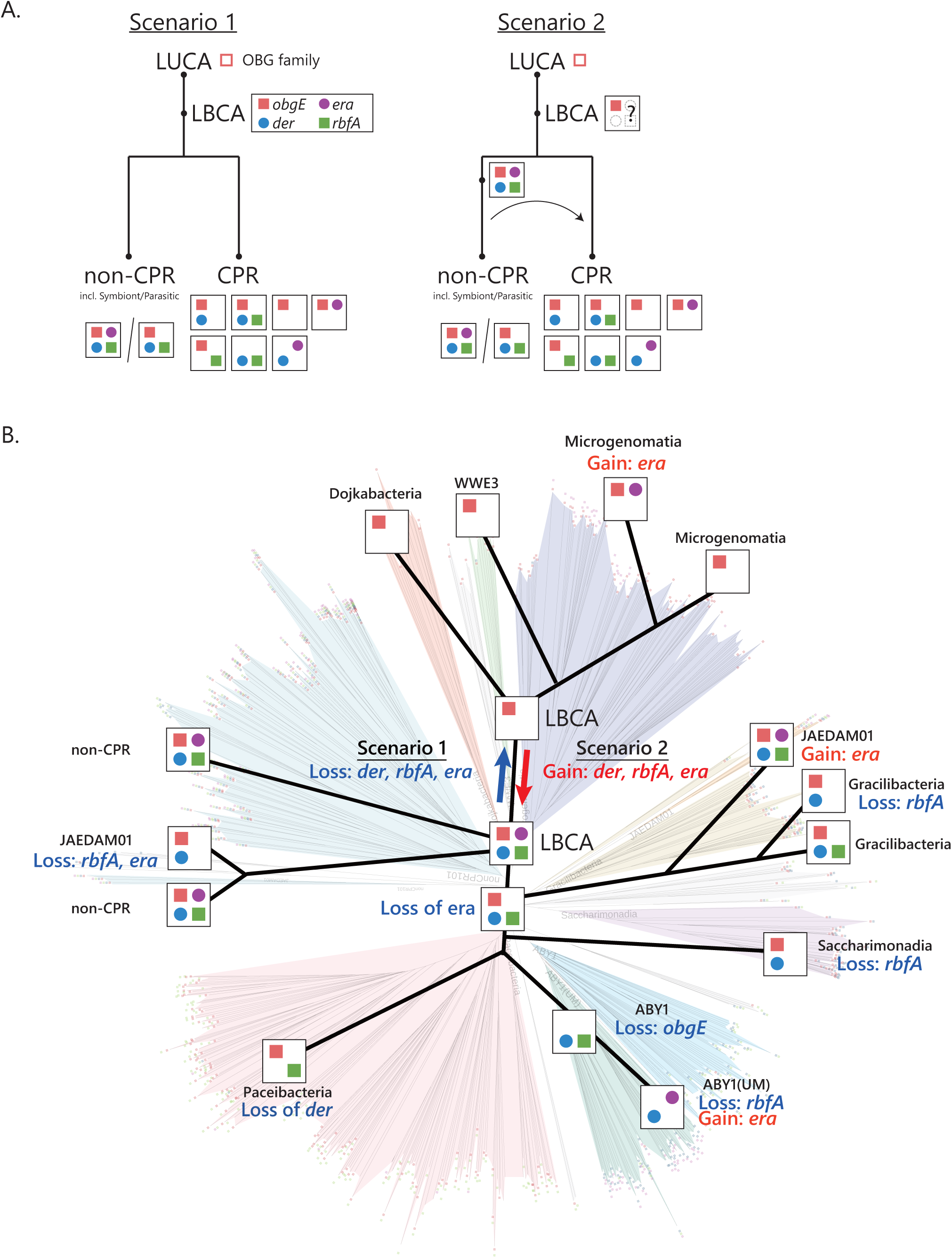
Evolutionary scenarios of ribosome biogenesis. (**A**) Schematic view of the evolutional history of ribosome biogenesis genes and their distribution among Candidate Phyla Radiation (CPR) and non-CPR from LUCA. (B) The tree illustrating the gain and loss events of four genes through evolution. The phylogenetic tree from Fig. 3A is displayed as a semi-transparent background layer beneath this tree.

## Discussion

The ribosome biogenesis process is generally considered to be conserved across all bacteria. However, our comprehensive genomic analyses of rRNA, RPs, and RBFs revealed that CPR bacteria possess a markedly reduced number of RBFs. This unique set of RBFs in CPR bacteria highlights the minimum requirements for RBFs in ribosome assembly and provides valuable insights into the evolution of ribosome biogenesis. Microorganisms with small cell and genome sizes, such as CPR and symbiotic/parasitic bacteria, tend to have reduced numbers of RBFs, suggesting that cell and/or genome size imposes constraints on the number of RBFs. Nevertheless, the repertoires of RBFs differ among these organisms, indicating that the composition of RBFs is presumably shaped by the evolutionary history of individual taxonomic linkage.

This study firstly provides evidence of co-evolution between ribosomal structures (rRNA and RPs) and ribosome biogenesis processes (Fig. 6). Although previous research has reported the co-evolution of rRNA and RP in chloroplastic, eukaryotic, and mitochondrial ribosomes (*28–32*), our comprehensive analyses demonstrate that *der* is the key factor in determining the secondary structure of rRNA and the presence of RPs in CPR bacteria. Although RBFs have been overlooked in discussions of ribosome evolution, they are likely important for the formation of ribosome structures throughout the evolutionary history of the ribosomes.

rRNAs, most RPs, and some RBFs have been used as molecular markers in the study of the evolution of life. Phylogenetic trees for individual RBF proteins, Der, ObgE, Era, and RbfA, align with the broader bacterial tree of life (Fig. 8). Moody et al. suggested that *der*, *obgE*, *era*, and *rbfA* are present in LBCA (*17*), whereas Leipe *et al.* proposed that the OBG family has been maintained since the emergence of the Last Universal Common Ancestor (LUCA) (*33*). Combining this information, one possible scenario is that the CPR bacteria lost the RBFs inherited from the LBCA, which included four RBFs (Fig. 9A, scenario 1). An alternative scenario is that the LBCA did not possess all four RBFs or only had one or two; however, organisms in the ecosystem during this evolutionary stage developed these RBFs, which were disseminated through gene transfer (Fig. 9A, scenario 2). CPR bacteria share similarities with symbiotic and parasitic bacteria in terms of genome size and the number of RBFs; however, symbiotic and parasitic bacteria typically encode four RBFs (Fig. 7D). Considering that RPs and rRNA co-evolved and that the ribosome is a macromolecular machine requiring a precise structure to function effectively, it is likely that the core components of the ribosome biosynthetic system cannot be drastically altered once established. In both scenarios, we assume that the current taxonomic classes of CPR bacteria and the ribosome were fundamentally established during the early stages of bacterial evolution.

Despite having fewer RBFs, CPR bacteria inhabit diverse terrestrial environments and constitute approximately 15% of the bacterial diversity, suggesting that they can maintain general translation activity even with their reduced ribosomal components. Although the mechanisms underlying the efficient ribosomal assembly in CPR bacteria remain unknown, several possible explanations exist. First, the ribosome structure in CPR bacteria differs slightly from that in non-CPR bacteria, which may lead to distinct conformational changes in the assembly intermediates (*34–36*). These differences may enable CPR bacteria to navigate the kinetic traps in the ribosome assembly process, even with fewer RBFs. Second, introns in the 16S and 23S rRNAs, which are rare in most bacteria but commonly found in CPR bacteria (*12*), may assist in forming the tertiary structures necessary for RP binding and proper rRNA folding. These introns are expected to be removed through self-splicing or endoribonuclease activity (*37*), potentially preventing mis-assembly or mis-folding. Third, there may be genes with yet-to-be-elucidated functions that contribute to ribosome assembly, compensating for the absence of certain RBFs. Testing these hypotheses based on studies on CPR bacteria should enhance our understanding of the functional adaptability of bacterial ribosome biogenesis. Moreover, in vitro ribosome reconstitution methods may provide valuable tools for evaluating these hypotheses (*38–41*).

In conclusion, although ribosome biogenesis was previously thought to be rigidly conserved across all bacteria, our study demonstrated that CPR bacteria exhibit potential flexibility and adaptability in their ribosome biogenesis. The unique characteristics of CPR bacteria demonstrate that even fundamental cellular processes such as ribosome biogenesis can exhibit substantial variation across different lineages. Future studies focusing on the molecular mechanisms underlying ribosome biogenesis in CPR bacteria could provide further insights into the evolutionary origins of cellular machinery and offer new perspectives on the diversity of life.

## Materials and Methods

### Dataset construction

Genome sequences for high-quality and non-redundant CPR bacteria were downloaded from the NCBI database. rRNA sequences were extracted using RNAmmer and barrnap (https://github.com/tseemann/barrnap), and protein sequences were predicted using Prodigal(*42*). Functional annotation of the predicted protein sequences was performed using the GhostKOALA and the KEGG Automatic Annotation Server (KAAS)(*43*, *44*). Subsequently, genomes containing all rRNA sequences with high completeness were selected using CheckM2(*20*). Several complete genomes of CPR bacteria, obtained from our metagenomic analysis, were initially processed using the Prokka in KBase platform and then same workflow as described above(*45*). For the analysis of non-CPR bacteria, annotation data were obtained from the ANNOTREE database(*19*), which includes information from over 30,000 genomes. From these, genomes registered in the RefSeq database were selected, and genome completeness was calculated using CheckM2.

### Phylogenetic analysis

Phylogenetic trees for *obgE*, *der*, and *era* were constructed using multiple sequence alignments of their GTPase domains, whereas the tree for *rbfA* was generated using full-length protein sequences. All phylogenetic analyses were performed with W-IQ-TREE under default parameters unless otherwise specified (*46*).

### Statistical analysis

PCA and factor loading analysis were performed by *prcomp* function in R(*47*).

The mutual information *MI(A,B)* has been calculated between genes A and B. Mutual information is maximum when there is complete co-variation in the occurrences of genes A and B, and it tends to zero as the co-variation decreases or the gene occurrences become independent. The calculations were as follows:

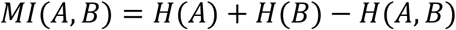

Here, *H(A)* represents the marginal entropy of the probability distribution *p(a)* of gene A’s abundance in CPR bacteria, summed over intervals in the probability distribution given by the formula: *H(A)* = -*Σp(a)ln p(a)*. Similarly, *H(A, B)* represents the relative entropy of the joint probability distribution *p(a,b)* of the occurrence of genes A and B across the CPR bacteria, summed over intervals in the joint probability distribution given by the formula: *H(A, B)* = -*ΣΣp(a, b)ln p(a, b)*. Using these values, the mutual information between genes A and B was calculated.

### Prediction of RNA secondary structures

Secondary structures of rRNA were predicted by Infernal’s *cmalign* using a covariance model constructed with the seed sequences from Rfam, a curated set of representative sequences for each family for the prediction of secondary structures of 5S rRNA(RF00001), 16S rRNA(RF00177), and 23S rRNA(RF02541) (*48*, *49*). The base number and region of the genes from *E. coli* were obtained from the Comparative RNA Web (CRW) site (*50*).

## Acknowledgments

We thank Dr. Tatsuhiko Hoshino and Dr. Hiroyuki Imachi for their support in constructing the CPR genome dataset, and Yuka Uenaka and Dr. Liao Fangyuan for their support in bioinformatics analyses at JAMSTEC.

## Author contributions

Conceptualization K.A. and S.S.; Methodology K.A. and S.I.; Investigation K.A., S.I., and S.S.; Visualization K.A.; Supervision Y.S. and S.S.; Writing original draft K.A., S.I., Y.S., and S.S.

---

**Supplementary Figure 1.** Boxplot representation of the number of RPs across different phyla in bacteria. Firmicutes divided into seven groups in the AnnoTree database.

**Supplementary Figure 2.** The combinations and the mutual information value. Small graph show different x-axis ranges.

**Supplementary Figure 3.** (**A**) The dependency network when the mutual information threshold is 0.05. (**B**) The dependency network when the mutual information threshold is 0.025.

**Supplementary Figure 4**. The boxplots represent the median length of helices. (A) 16S rRNA (B) 5S rRNA (C) 23S rRNA.

**Supplementary Figure 5.** (**A**) Tertiary structure of L1 stalk (Silver), H78 (Purple), uL1 (Blue), and Der (Yellow). (PDB: 3J8G)

**Supplementary Figure 6.** (**A**) The lengths of H78 length in CPR bacteria are arranged in order of length. **(B)** in Paceibacteria, ABY1, and ABY1(UM) **(C)** in Dojkabacteria, WWE3, and Microgenomatia **(D)** in JAEDAM01, Gracilibacteria, and Saccharimonadia.

**Supplementary Figure 7. Boxplot representation of the genome size across different phyla**. (**A**) Genomes in the AnnoTree with the completeness of 95 or more and registered for the RefSeq were selected. The red circle represents the genome size of symbiotic bacteria species. (**B**) Colors of the plots in Figure 7C were modified, with the CPR and top six classes containing the highest number of species displayed in the legend.

**Supplementaly Table 1**.

This table presents the curated list of RBFs used in this study. KEGG_ID refers to the KEGG orthology ID. Name indicates the commonly used name of the factor, while Synonym lists any alternative names. Category in this study describes the classification or role of each factor based on the current research. Category in KEGG Pathway (Ribosome biogenesis in prokaryotic type) refers to the corresponding category in the KEGG database. Essentiality (PEC) and Essentiality (KEIO) columns reflect the factor’s essentiality as defined by the Profiling of E.coli Chromosome (PEC, https://shigen.nig.ac.jp/ecoli/pec/) and the KEIO collection, respectively(*8*, *51*). BAC120 and LBCA indicate the presence of the factors in the BAC120 phylogenetic marker set and in the last bacterial common ancestor (LBCA), respectively(*11*, *52*). References provide key literature for RBFs that isn’t included in the KEGG database, and the Note column contains any additional relevant annotations.

**Supplementary Table 2.**

This table presents ribosome biogenesis factors and their conservation across CPR and Non-CPR bacterial lineages. KEGG_ID refers to the KEGG orthology ID associated with each factor. symbol denotes the gene symbol commonly used for each factor. Category describes the functional classification. non-CPR (%) and CPR (%) indicate the conservation of each factor within Non-CPR and CPR bacteria, respectively.

**Supplementary Table 3.**

This table displays the mutual information (MI) values for gene pairs (Gene A and Gene B) and the distribution of species based on the presence or absence of these genes in CPR bacteria.

**Supplementary Table 4.**

This set of tables presents the number of gene interactions (edges) and the total mutual information (MI) for each gene, calculated at three different MI thresholds: greater than 0.1, 0.05, and 0.025. The genes are ranked in descending order based on their total MI values for each threshold.

## Supplementary Information

### Supplementary_information_CPR_505.csv

This table provides detailed information on CPR genomes. Each column is described as follows. Assembly Accession: The unique accession number assigned to each genome assembly. Organism Name: The taxonomic name of the organism based on genome data. Rep. genome: Indicates whether the genome is a complete representative genome (Comp) or partial. Sum of tRNAs: The total number of tRNA genes identified in each genome. Completeness: The estimated completeness by CheckM2 of each genome, expressed as a percentage. Contamination: The estimated level of contamination by CheckM2 in each genome, expressed as a percentage. Size: Genome size in megabases (Mb). GC: GC content percentage of the genome. Scaffolds: The number of scaffolds in each genome assembly. WGS: The Whole Genome Shotgun (WGS) project identifier. ncbi_organism_name: The organism name as recorded in the NCBI database. ncbi_taxonomy: The taxonomy classification according to NCBI. gtdb_taxonomy: The taxonomy classification according to the Genome Taxonomy Database (GTDB). GTDB_Phylum: The phylum classification based on GTDB taxonomy. GTDB_Class: The class classification based on GTDB taxonomy. Group: Classification group of the organism in this study

### Supplementary_information_nonCPR.csv

This table provides detailed information on non-CPR genomes. Each column is described as follows. Assembly Accession: The unique accession number assigned to each genome assembly. Organism Name: The taxonomic name of the organism based on genome data. nonCPR107: Indicates whether the genome is part of nonCPR107 as shown in Figure 3A.

### 23S_rRNA_aln_L1stalk.fasta

Alignment of the 23S rRNA seed sequences from Rfam with the 23S rRNA sequences of CPR bacteria, with regions corresponding to helices H76, H77, and H78 extracted and presented in FASTA format.

