## Supplementary Figures for "Evolutionary Flexibility of Ribosome Biogenesis in Bacteria"

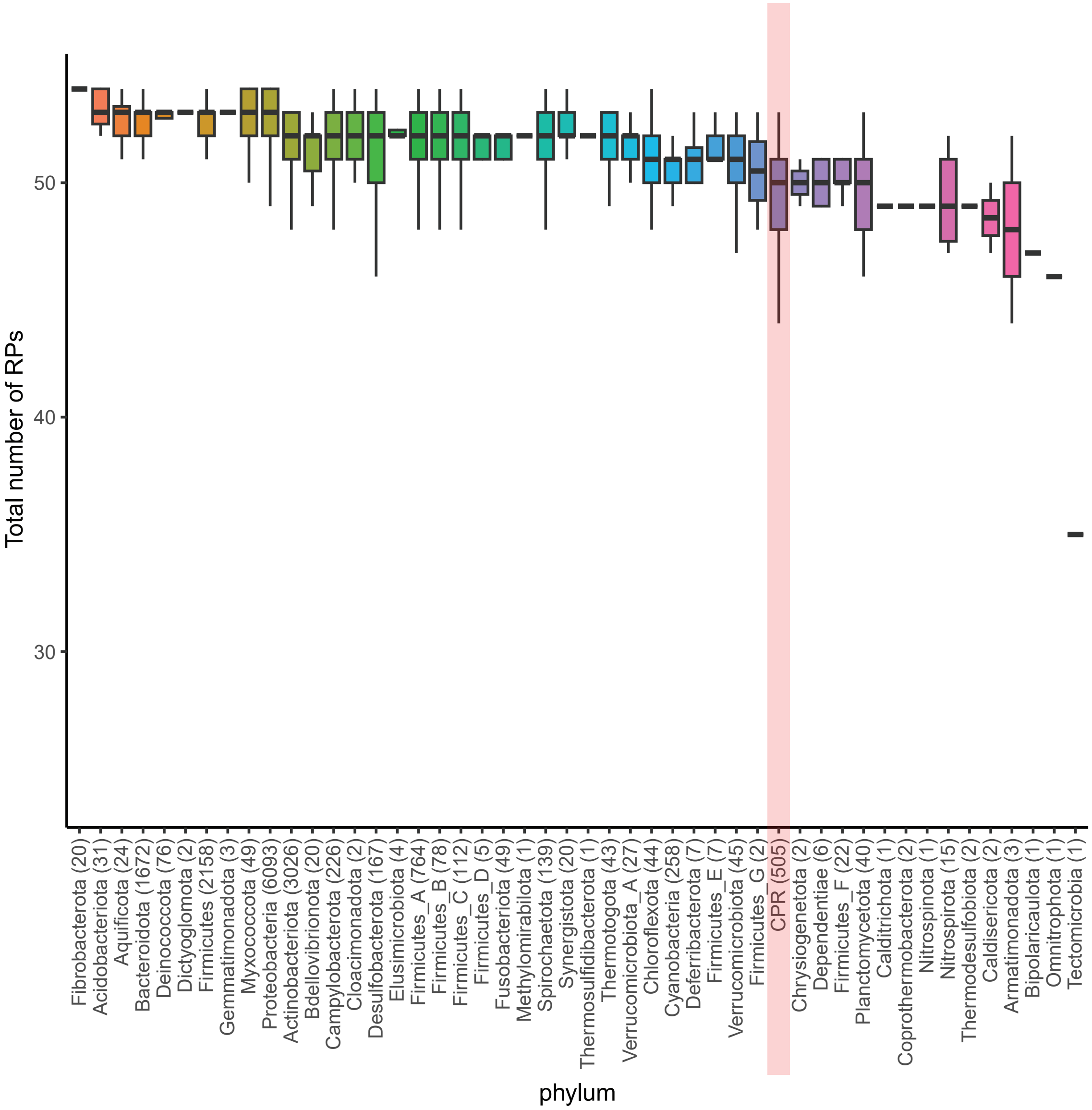


**Supplementary Figure 1**

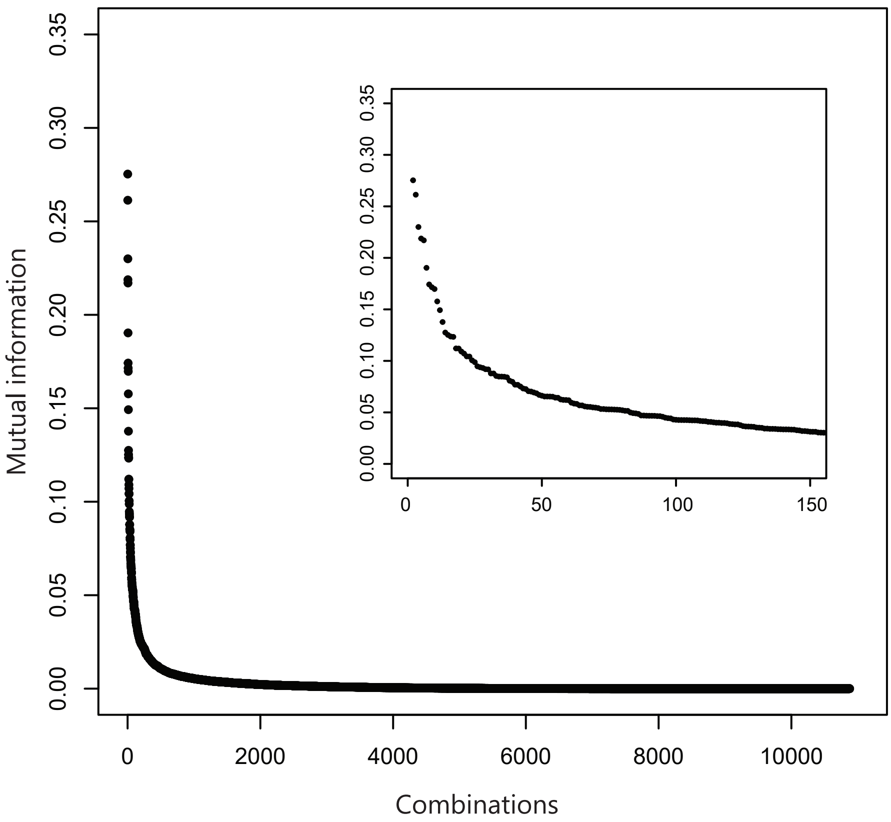


**Supplementary Figure 2**

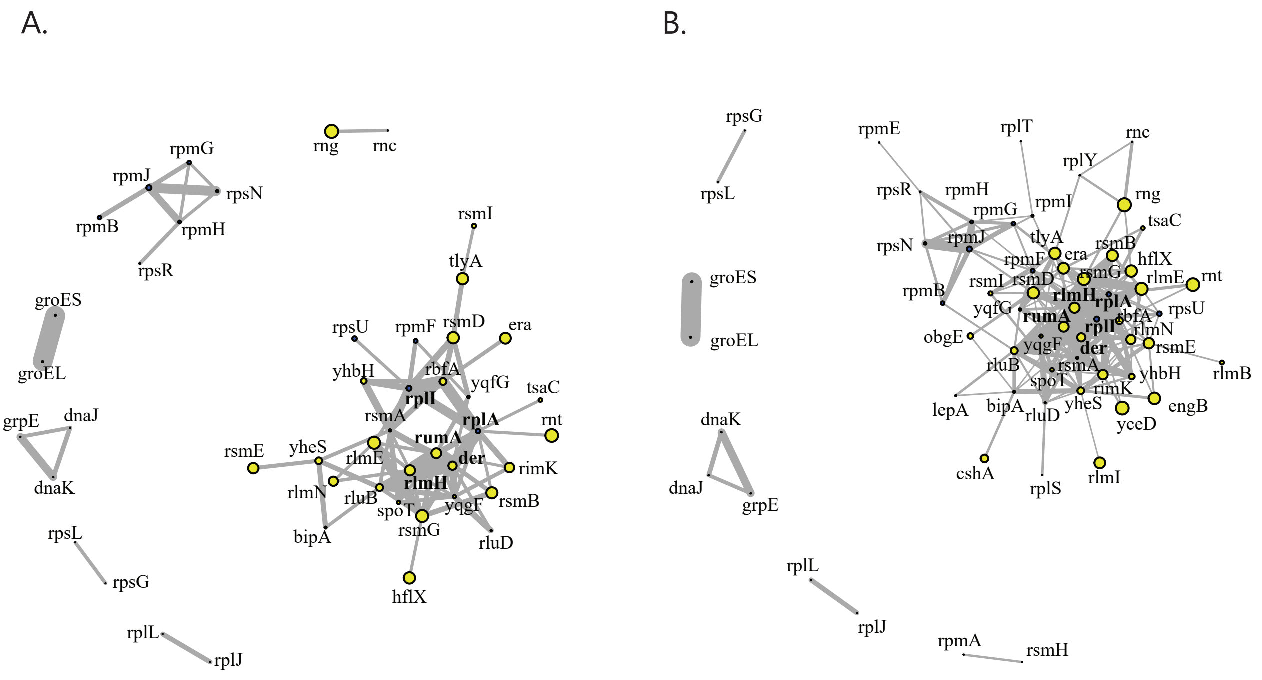


**Supplementary Figure 3**

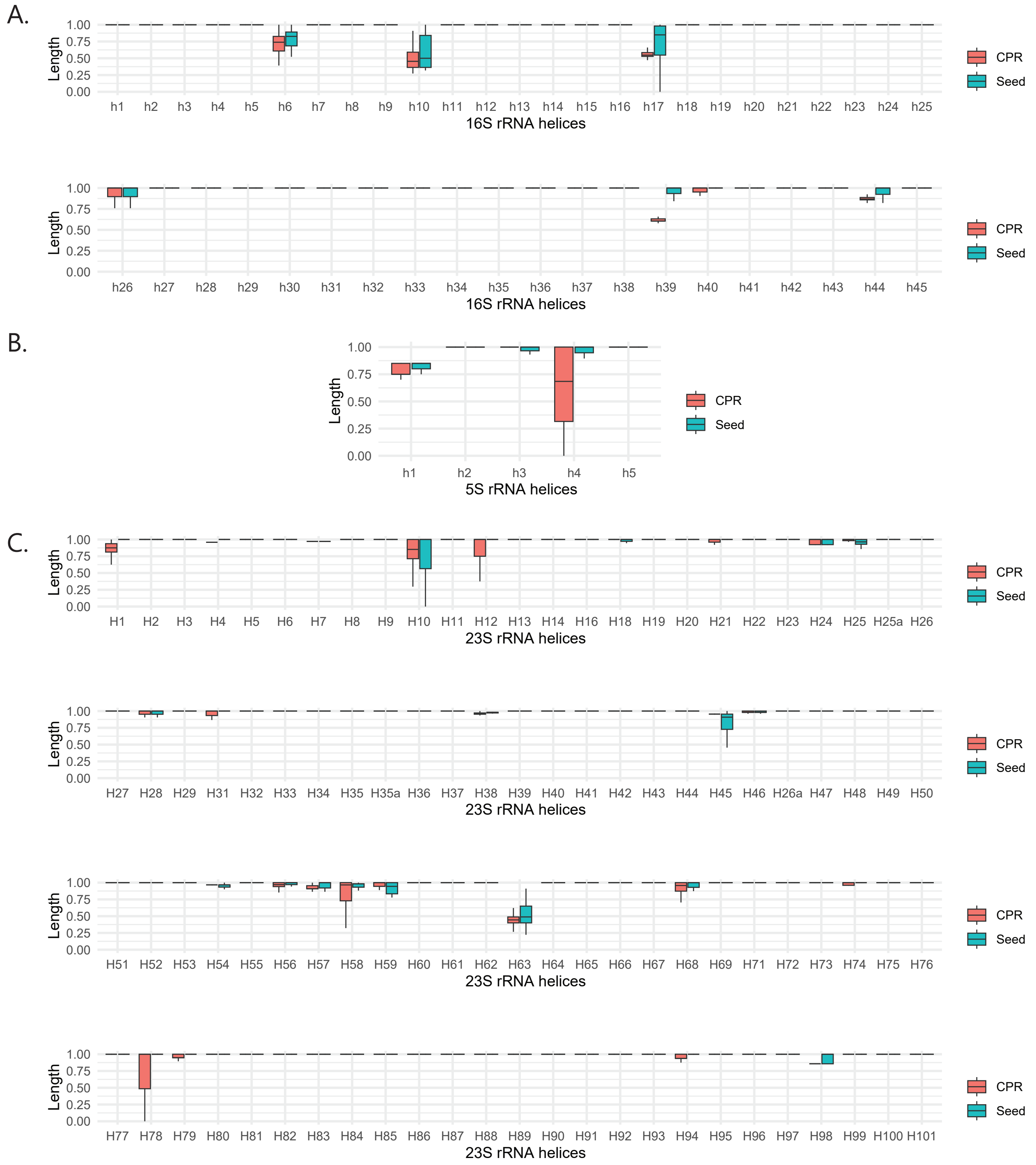


Supplementary Figure 4

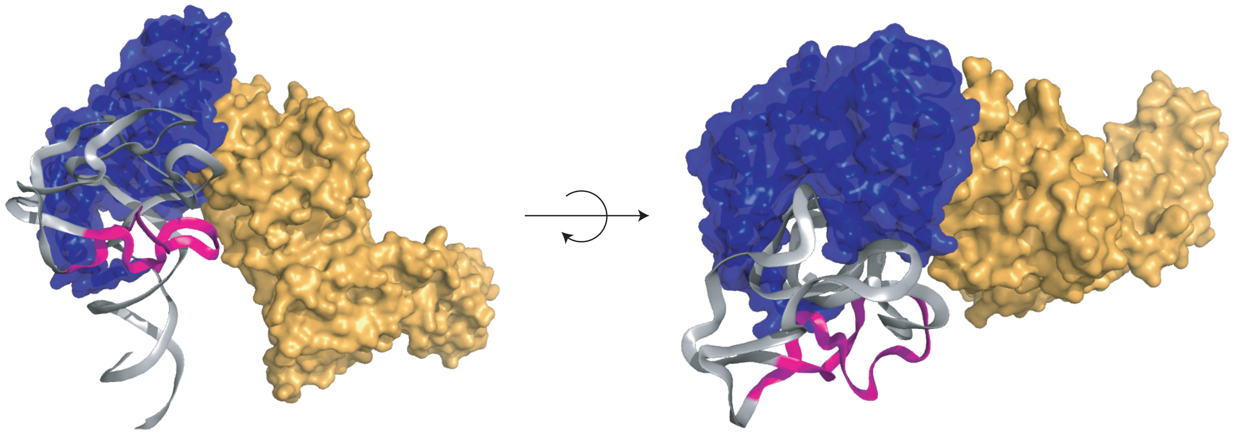

**Supplementary Figure 5**

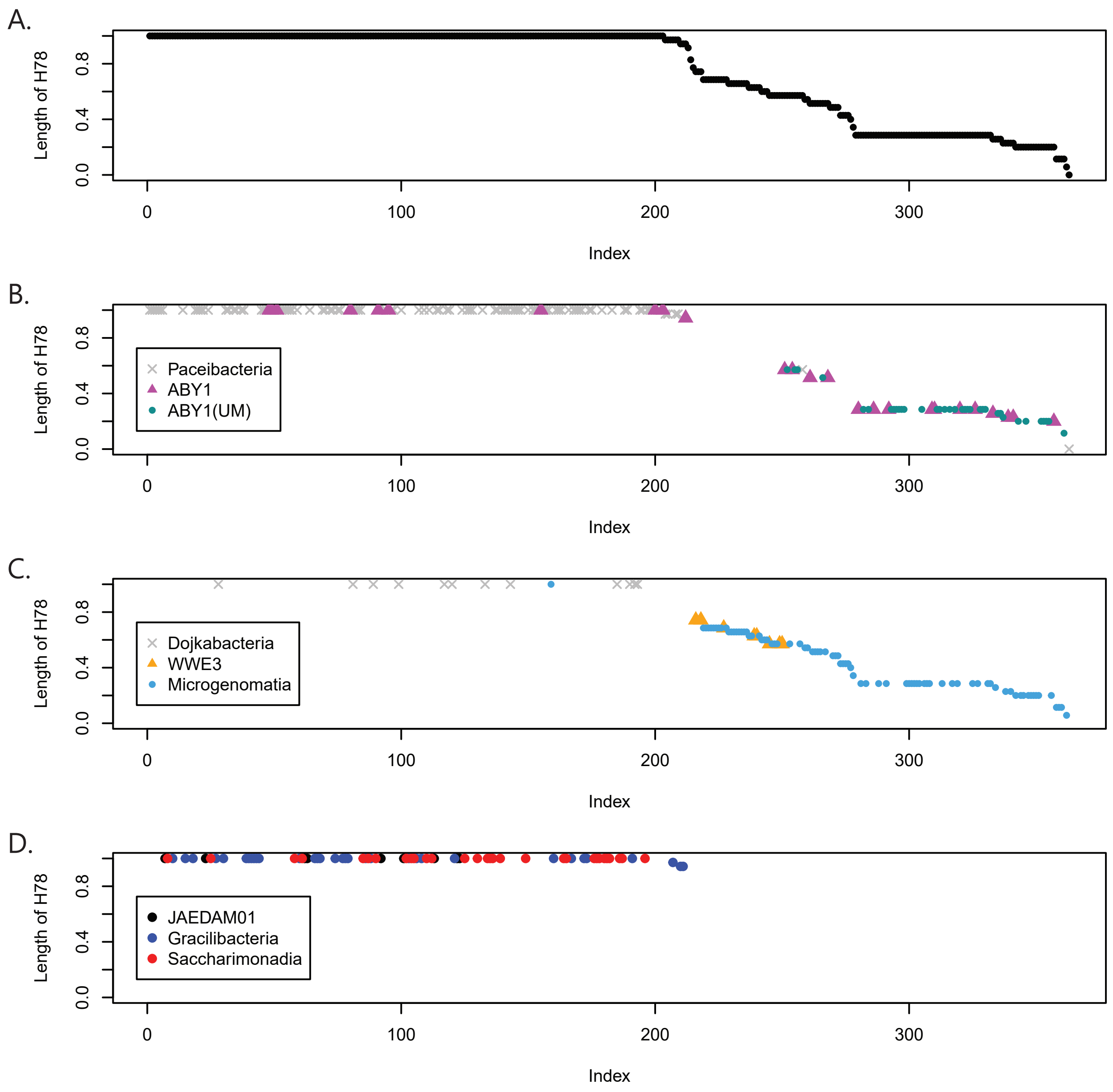


Supplementary Figure 6

A.

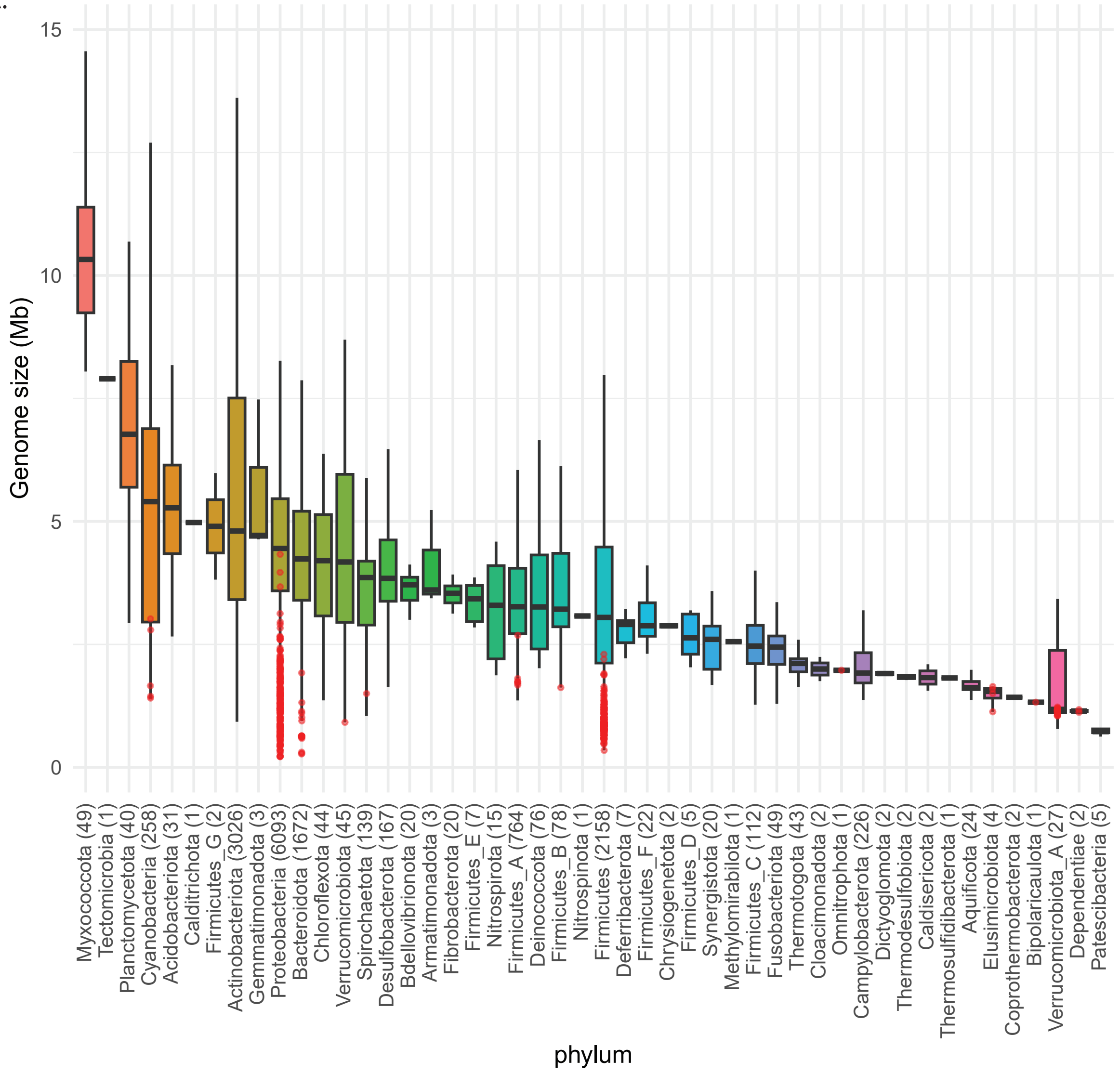


B.

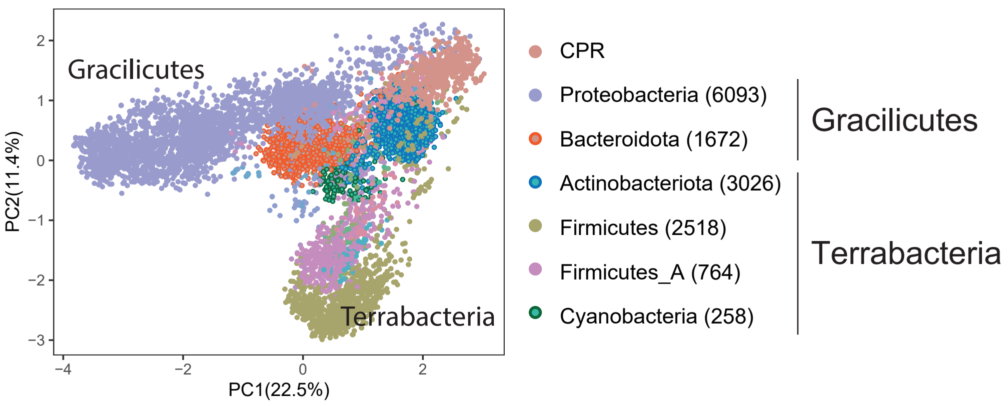


**Supplementary Figure 7**
